# Pathway-Based Analysis of Metabolic Enzymes as Drug Targets

**DOI:** 10.64898/2026.10.21.754949

**Authors:** Deepti Subray Bhat, Bhupender Singh

## Abstract

Tuberculosis (TB), caused by Mycobacterium tuberculosis, remains a major global health crisis, aggravated by rapid emergence of multidrug-resistant (MDR) and extensively drug resistant (XDR) strains. Conventional drug discovery is time-consuming and expensive, making computational target identification and drug repurposing as crucial strategies to accelerate the development of novel anti-tubercular therapies. This study aimed to identify and prioritize the metabolic enzymes as potential drug targets through pathway based analysis and to screen the FDA-approved drug library for potential canidates for drug repurposing. Based on the pathway analysis, **para-aminobenzoate** synthase component I (PabB) was selected as a potential drug target owing to its role in folate biosynthesis. Subesquently, an FDA-approved drug library was screened based on ADMET properties and selected compounds were used for molecular docking agaist the protein. Molecular docking through DiffDock and validation using GNINA revealed that Ligand 5 and Ligand 27 exhibit favourable interaction profiles within the binding region of PabB, and emerged as potential candidates for further investigation in drug repurposing. The identification of PabB protein as an essential metabolic drug target, coupled with screening of FDA-approved drugs as inhibitors, provides a promising framework for anti tubercular drug discovery.

## Introduction

Tuberculosis(TB) is a communicable disease caused by *Mycobacterium tuberculosis* which primarily attacks lungs but can also affect other parts of the body. It affects millions of people globally(Waghapure et al., n.d.). ***M. tuberculosis*** has the ability to remain dormant within the host and develop resistance to anti-tubercular drugs. To overcome this new anti-tubercular agents are actively being searched against multidrug-(MDR) and extensively drug-resistant (XDR) *M. tuberculosis*. Efforts are being made to identify novel drug targets(Saxena and Singh 2019a). The computational approach has been used to investigate novel drug targets in other pathogenic organisms such as *Pseudomonas aeruginosa* and in *Helicobacter pylori* (Saxena and Singh 2019b),(Akter et al. 2026). The identification of essential genes provides a systematic approach for exploring proteins that are indispensable for the survival of the pathogenic organisms (Amir et al. 2014). Despite these extensive research carried out on tuberculosis and the availability of several anti-tubercular drugs, there is still a need to identify and validate new, pathogen specific targets. Computational approaches that combine essential gene analysis, functional annotation, pathway analysis, and protein interaction studies can help identifying and prioritizing promising drug targets in *M. tuberculosis*(Alanzi et al. 2025),(Syed et al. 2025). In the present study, essential proteins of *M. tuberculosis* were analysed using a computational target identification approach. The protein sequences were retrieved and screened based on their similarity to human proteins to identify potential targets with minimal similarity to the host. A suitable candidate was then selected for further functional and pathway characterization and consider for further investigation as a potential drug target for anti-tubercular drug discovery. While identification of potential drug targets is critical step, subsequent discovery and development of new drugs remains a lengthy and resource intensive process. Drug repurposing offers an alternative strategy by investigating existing drugs for new therapeutic applications(Kulkarni et al. 2023),(Saranraj and Kiran 2025). FDA-approved drugs are particularly attractive for the identification of potential drug candidates as these have already undergone substantial pharmacological safety and evaluation. Therefore, in the present study FDA-approved drugs were explored as potential candidates against selected *M. tuberculosis* target using ADMET based filtering following molecular docking.

## Methods

Essential genes of *M. tuberculosis* and their corresponding protein sequences in FASTA format were retrieved from Database of Essential Genes(DEG) (http://tubic.tju.edu.cn/deg/) (Zhang et al. 2004). The retrieved essential protein sequences of *M. tuberculosis* were used as a query for running local BLASTP(Basic Local Alignment Search Tool for Proteins) against human proteome using **ncbi-blast-2.17.0+**.The local BLASTP results were filtered to identify proteins homologous to human proteins based on an E-value of <10^-4^ and sequence identity of >35%. Proteins meeting both the criteria were considered homologous and were excluded from further analysis. The remaining proteins were considered non homologous and were selected for subsequent analysis (Dar et al. 2020),(Ayu Eka Pitaloka et al. 2023).One of the non-homologous proteins was selected as a potential drug target based on its relatively higher E-value and lower percentage identity to the human proteins.The protein structure and associated functional annotations were retrieved from **UniProtKB** database (https://www.uniprot.org/uniprotkb/O05591/entry). KEGG was used to identify the corresponding KEGG Orthology(KO) assignment and to identify the metabolic pathways associated with the protein (https://www.genome.jp/kegg/). **QuickGO** was used to perform Gene Ontology (GO) annotation to characterize the molecular function, biological process, and cellular localization of the selected protein(https://www.ebi.ac.uk/QuickGO/). **STRING(Search Tool for the Retrieval of Interacting Genes/Proteins)** was used to analyse the protein-protein interaction network and identify functionally associated proteins and biological pathways (https://string-db.org/).**Therapeutic Target Database** was used to check the presence of the protein as an already established target.**FDA-approved drugs’ dataset** was obtained using https://enamine.net/compound-libraries/bioactive-libraries/fda-approved-drugs-collection. **ADMET-AI** (https://github.com/swansonk14/admet_ai) which was installed and executed locally was used for ADMET based filtering of FDA-approved drugs. The predicted physicochemical, drug-likeness and toxicity-related properties were subsequently used for further filtering according to predefined criteria.Candidate selection for molecular docking was done by retaining the drugs based on overall predicted and favourable Drug likeness and ADMET profiles.The selected candidate drugs were subjected to molecular docking against PabB protein using **DiffDock** at https://build.nvidia.com/mit/diffdock (Corso et al. 2023) to obtain their confidence scores and predict their binding poses.The top-ranked docking poses were validated using **GNINA** at https://github.com/gnina/gnina (McNutt et al. 2025). The resulting Minimized affinity, CNN score, CNN affinity were recorded. Based on the docking and validation results, four Ligand-PabB complexes were selected for further protein-ligand interaction analysis using the Protein-Ligand Interaction Profiler at https://plip-tool.biotec.tu-dresden.de/plip-web/plip/index (“PLIP 2025: Introducing Protein–Protein Interactions to the Protein–Ligand Interaction Profiler | Nucleic Acids Research | Oxford Academic,” n.d.). The identification of amino acid residues involved in interaction with the protein were recorded.The protein-ligand complexes were visualized using **PyMOL** (https://www.pymol.org/).

## Results

### Identification of essential genes and retrieval of protein sequences in FASTA format

A total of 614 genes were obtained from DEG database (https://tubic.org/deg/public/index.php/organism/bacteria/DEG1010.html)

The FASTA format of all those protein sequences were obtained through DEG database (http://tubic.org/deg/public/download/DEG10.aa.gz).

### Local BLASTP and identification of non-homologous proteins

The retrieved essential protein sequences subjected to BLASTP analysis against the human proteome to identify proteins with significant similarity to human proteins and identify non-homologous proteins.

### Selection of the potential drug target

Among non-homologous proteins **para-aminobenzoate synthase component I (Pab B)** (https://tubic.org/deg/public/index.php/information/bacteria/DEG10100150.html) was chosen as a potential drug target candidate based on its low sequence identity (28.743%) and high E-value of 4.3.

### UniprotKB results

The protein **para-aminobenzoate synthase component I (PabB)** was characterized using UniProtKB. The database provided information regarding the protein"s functional annotation, sequence and alpha fold structure

### KEGG pathway analysis

The DEG entry associated with the selected protein contained multiple KEGG Orthology (KO) identifiers. However, the corresponding UniProtKB entry assigned a single KO identifier to the selected protein.

The selected protein was mapped to KEGG Orthology (KO) identifiers in KEGG mapper tool (https://www.genome.jp/kegg/mapper/search.html) and the associated pathways are identified.

K01665 was identified as the KEGG Orthology (KO) identifier for pabB gene, which encodes para-aminobenzoate synthase component I. KEGG associates K01665 with the folate biosynthesis and biosynthesis of cofactors.

Folate metabolism is important for one-carbon transfer and nucleotide/amino-acid biosynthesis. The pathway contains enzymes that can potentially be exploited for drug targeting (Thiede et al. 2016)

PabB (K01665), encoding **para-aminobenzoate synthase component I (PabB)**, is involved in the biosynthesis of 4-amino-4-deoxychorismate, an intermediate in PABA biosynthesis. According to KEGG, the associated reaction (R01716) converts chorismate and L-glutamine into 4-amino-4-deoxychorismate and L-glutamate. This reaction is part of the pathway leading to PABA, which subsequently contributes to folate biosynthesis.

### QuickGO analysis

Gene ontology annotations were obtained using QuickGO. The annotations support the role of para-aminobenzoate synthase component I in folate biosynthesis.(https://www.ebi.ac.uk/QuickGO/annotations?geneProductId=O05591)

### STRING analysis

The protein-protein interaction network of para-aminobenzoate synthase component I (PabB) from *M. tuberculosis* was analysed using STRING database (https://string-db.org/network/83332.Rv1005c).This analysis showed that the interacting proteins were predominantly involved in aromatic amino acid biosynthesis and tryptophan biosynthesis. The other processes are listed in the Figure 6. This supports the functional importance of ‘pabB’ gene and the protein encoding the same in *M. tuberculosis*.

**Figure 1.**
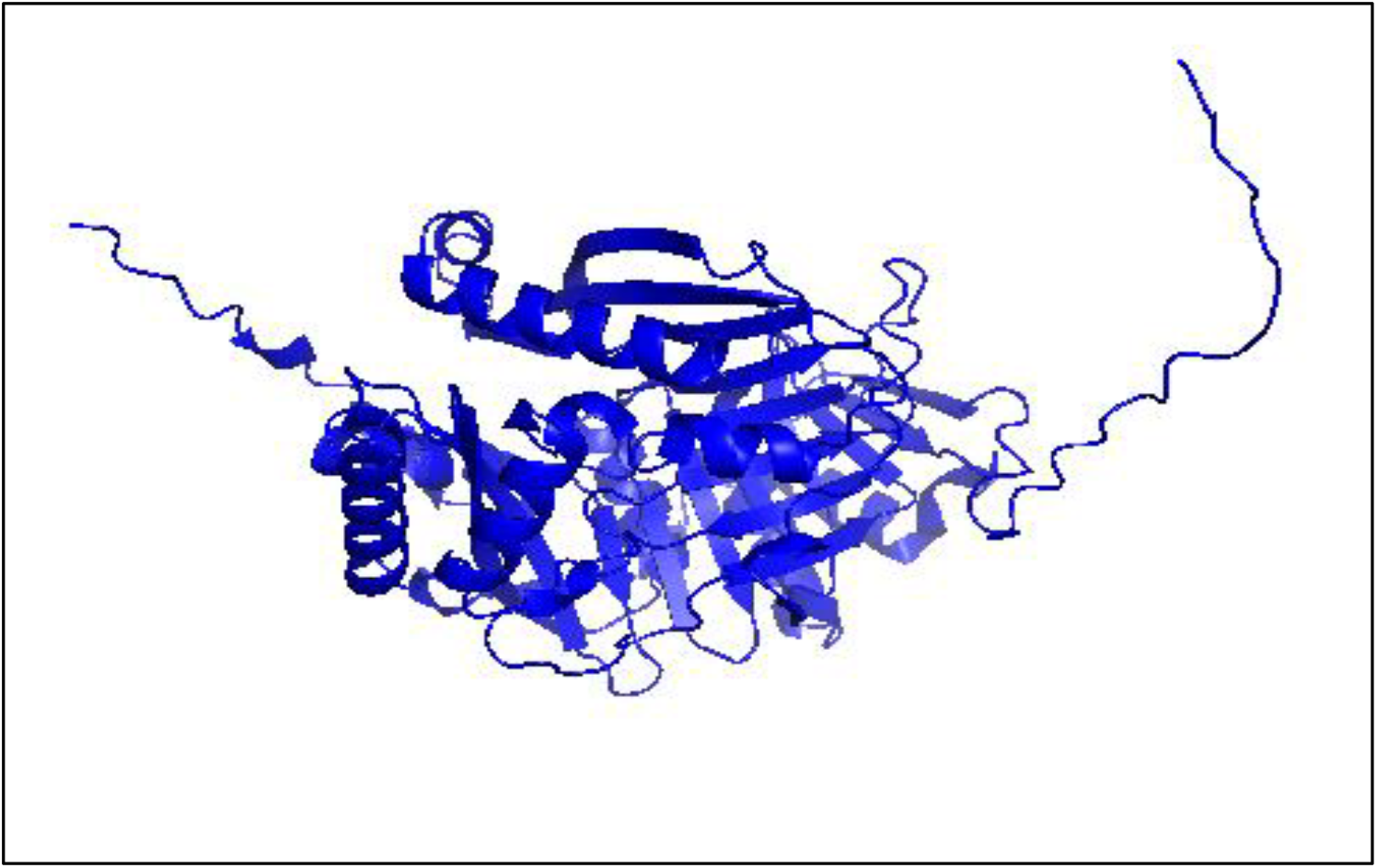
AlphaFold structure of para-aminobenzoate synthase component I (PabB).

**Figure 2.**
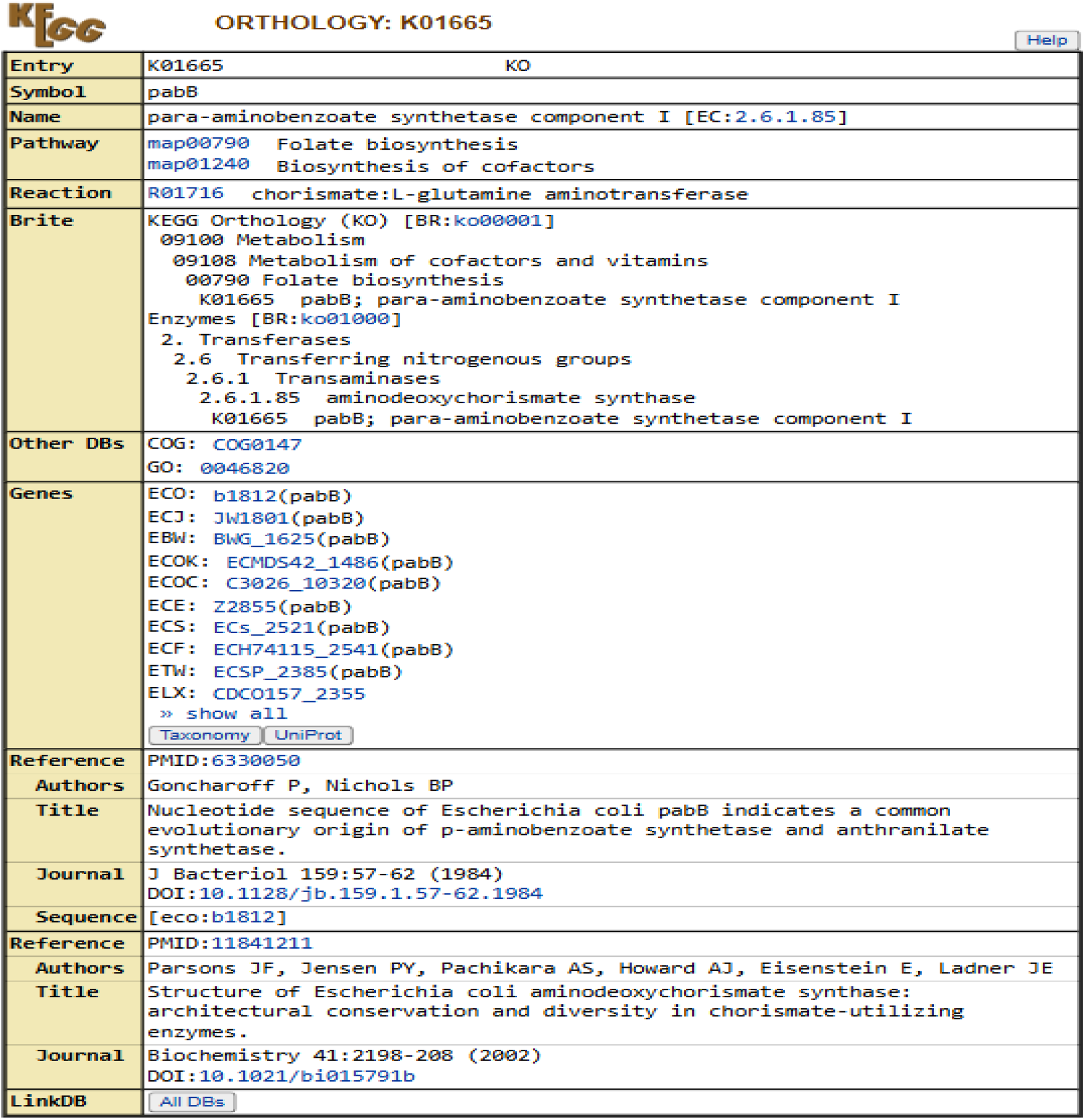
KEGG Orthology of para-aminobenzoate synthase component I (PabB)

**Figure 3.**
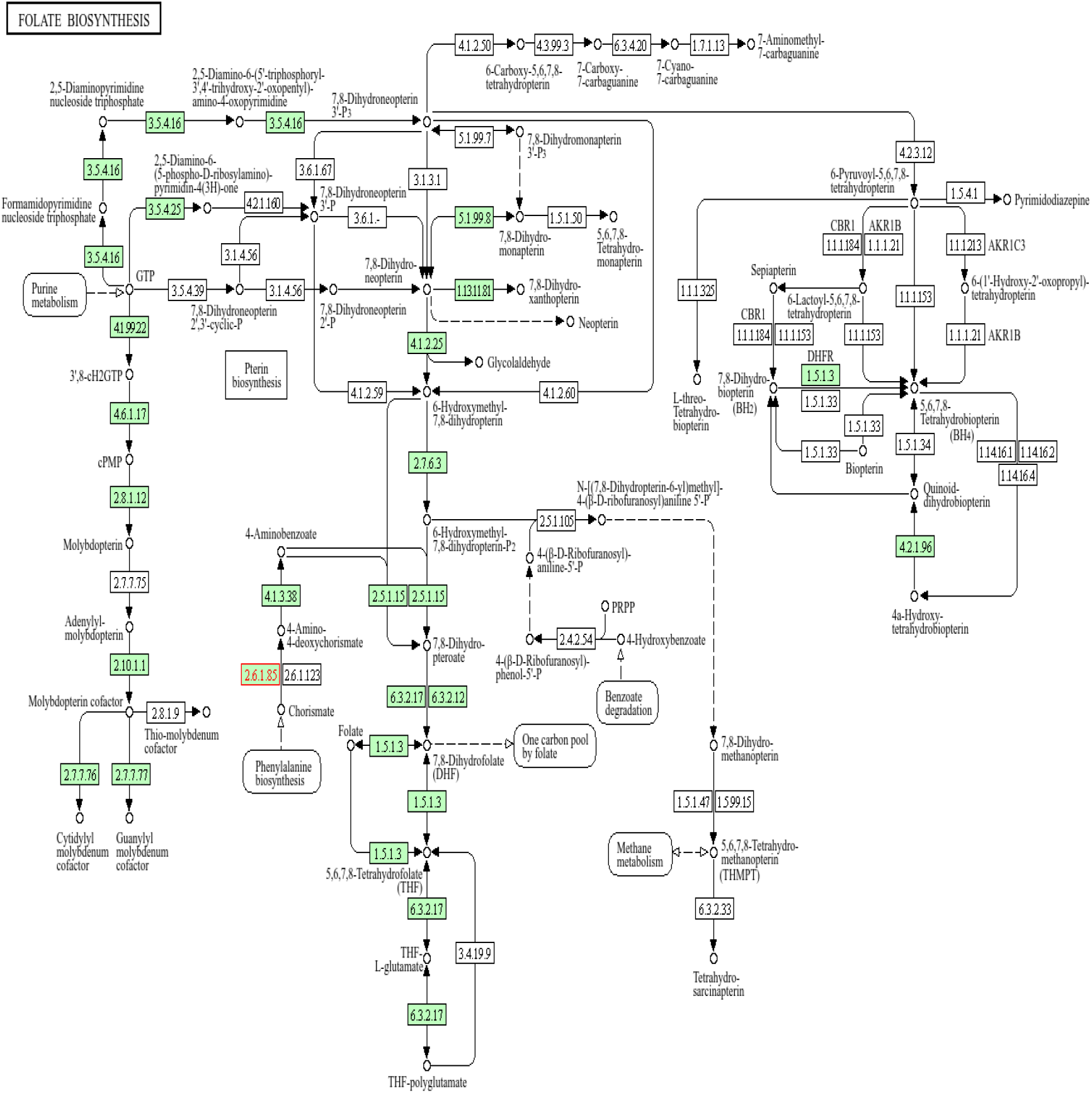
KEGG folate biosynthesis pathway of M. tuberculosis.

**Figure 4.**
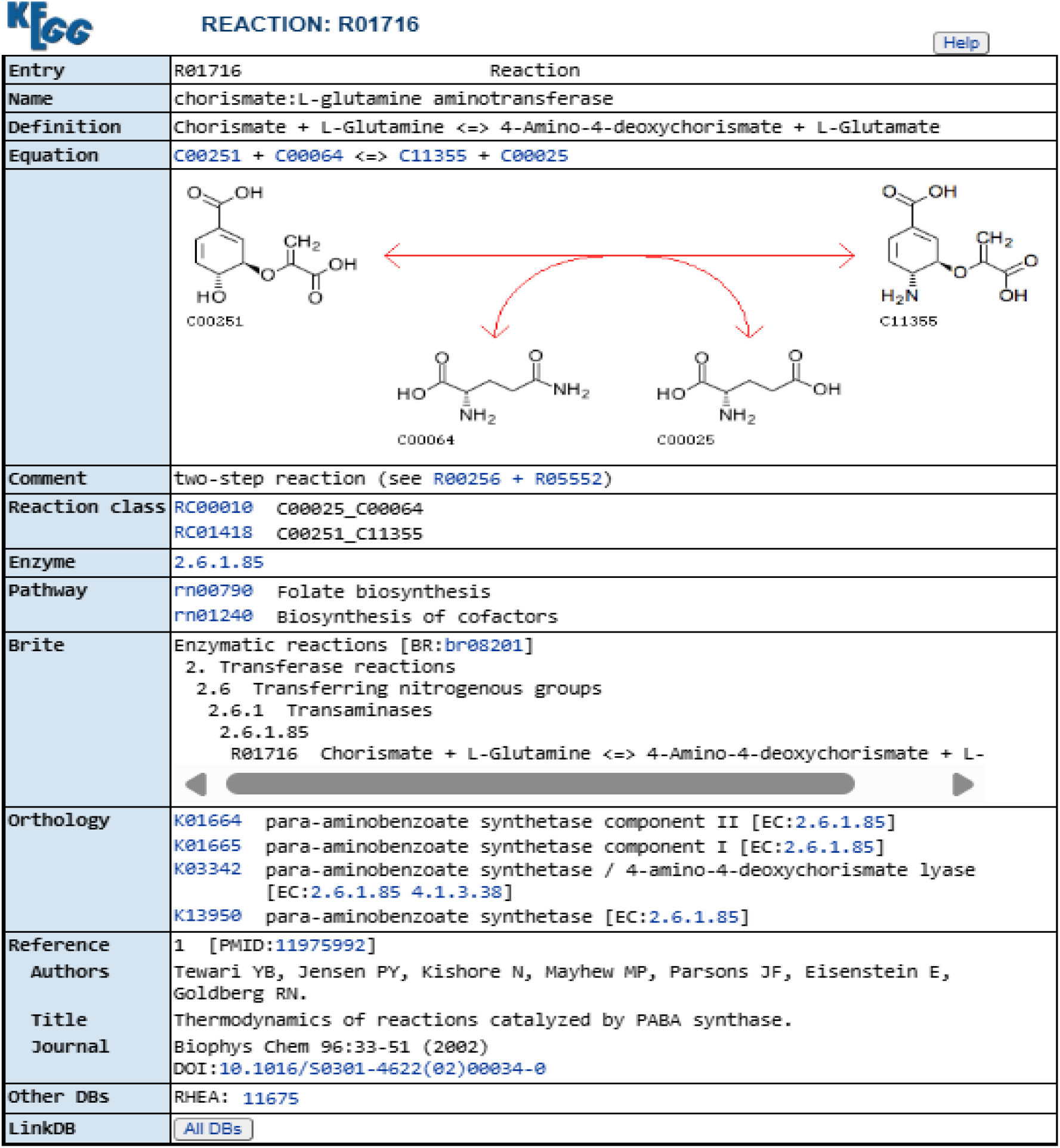
Reaction catalyzed by para-aminobenzoate synthase component I (PabB).

**Figure 5.**
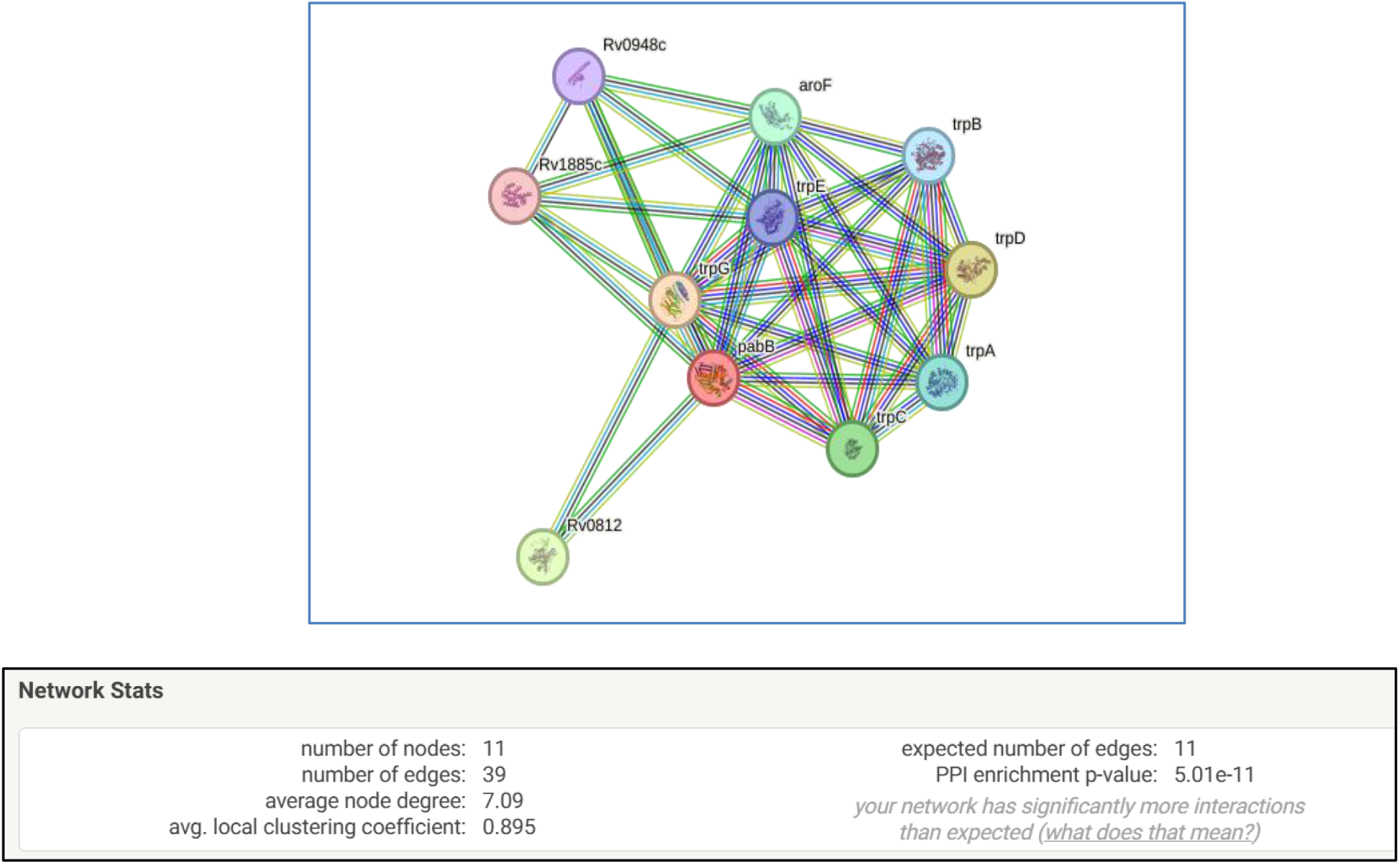
STRING analysis of para-aminobenzoate synthase component I(PabB)

**Figure 6.**
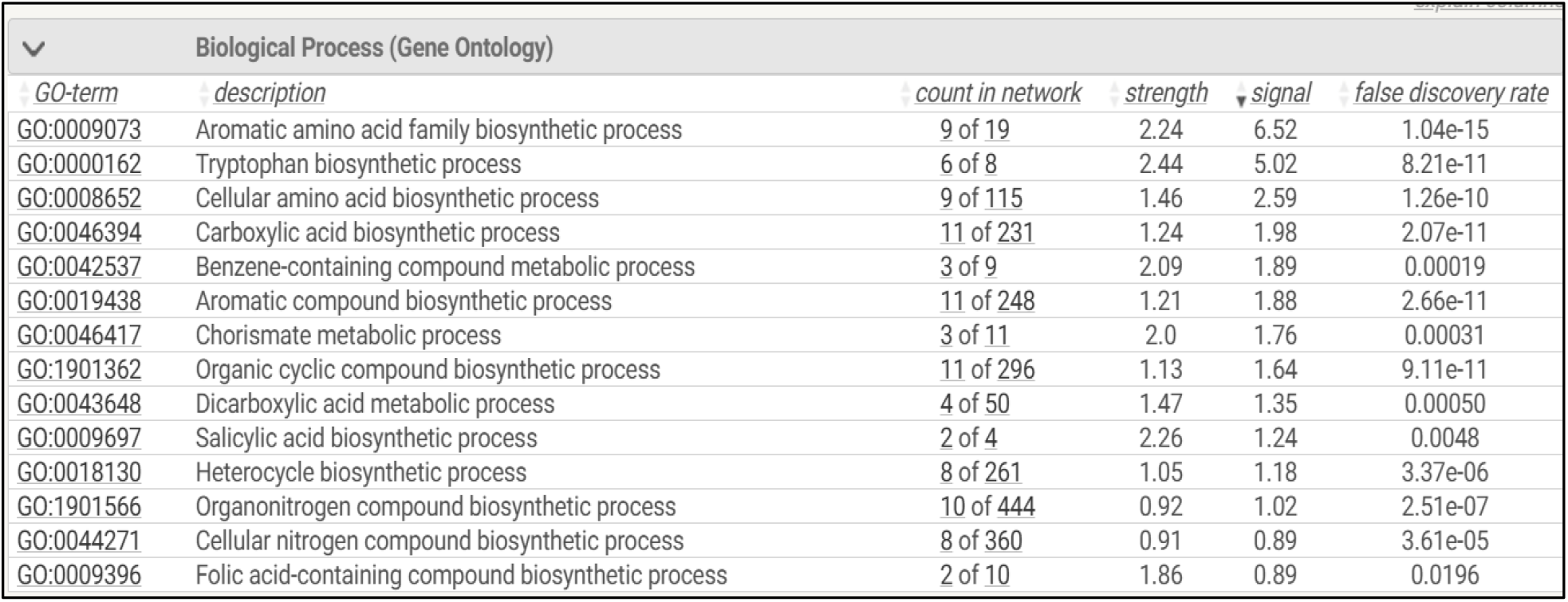
Biological Processes associated with para-aminobenzoate synthase component I (PabB) from STRING analysis.

### Therapeutic Target Database (TTD) analysis

No corresponding entry for para-aminobenzoate synthase component I (PabB) of *M. tuberculosis* is listed in the TTD (https://db.idrblab.net/ttd/) indicating that it may represent a relatively unexplored target and it is selected for further analysis.

### Retrieval of FDA-approved drugs

A total of 1123 FDA-approved drugs were retrieved from https://enamine.net/compound-libraries/bioactive-libraries/fda-approved-drugs-collection. They were further subjected to ADMET based filtering.

### ADMET-AI analysis

ADMET-AI based screening resulted in the 59 candidate compounds meeting the applied filtering criteria.("Ligand 1-59"were study-specific labels assigned to these 59 candidate compounds and the Catalog ID refers to the compound identification number assigned by the enamine database).The chemical name of the shortlisted compounds were retrieved from PubChem database (https://pubchem.ncbi.nlm.nih.gov/) by performing SMILES based search .From these compounds, 10 candidates were prioritized for molecular docking against para-aminobenzoate synthase component I (PabB) by using a multi parameter assessment incorporating drug-likeness, QED, solubility, HIA, Caco-2, AMES, DILI, carcinogenicity and hERG.

### DiffDock and GNINA analysi

The 10 selected ligands were taken for molecular docking using DiffDock. Two of the ten selected ligands (Ligand 45 and Ligand 48) showed parsing error and were excluded.The remaining ligands were docked against PabB protein using DiffDock and the resulting DiffDock confidence scores were recorded for top-ranking pose for each ligand. The results of DiffDock were validated using GNINA docking tool and the corresponding **GNINA Minimized Affinity(kcal/mol), GNINA CNN Score, GNINA CNN Affinity**.Based on the overall docking and validation results, Ligands 1, 5, 20 and 27 were selected for subsequent protein-ligand interaction analysis using **PLIP**.

### PLIP Analysis

The visualizations of the docked conformations of the selected 4 ligands against PabB are shown in the Figure 7. The amino acid residues interacting with the protein and the type of interactions are listed in the Table 9.Both hydrophobic and hydrogen bond interactions were observed in all of the four docked conformations.From the docked conformations, it is visible that VAL385 and ASN397 residues of PabB interact with all of the four ligands which can be predicted as they interact with a common region of PabB suggesting a conserved interaction region within the predicted binding site.

**Table 1.** Results of Local BLASTP of *M. tuberculosis* essential proteins against human proteome.

| Description | Number of proteins |
| --- | --- |
| Essential *M.tuberculosis* protein sequences retrieved from DEG | 614 |
| Human proteome sequences used for BLASTP analysis | 147506 |
| Proteins showing homology with human proteins | 154 |
| Non-homologous proteins obtained after exclusion | 460 |

**Table 2.** BLASTP Parameters of para-aminobenzoate synthase component I (PabB).

| Query ID | Subject ID | Percentage identity | Alignment length | E-value | Bit score |
| --- | --- | --- | --- | --- | --- |
| DEG10100150 | tr\|A0ACI8VK40\|A0ACI8VK40_HUMAN | 28.743 | 167 | 4.3 | 32.3 |

**Table 3.**
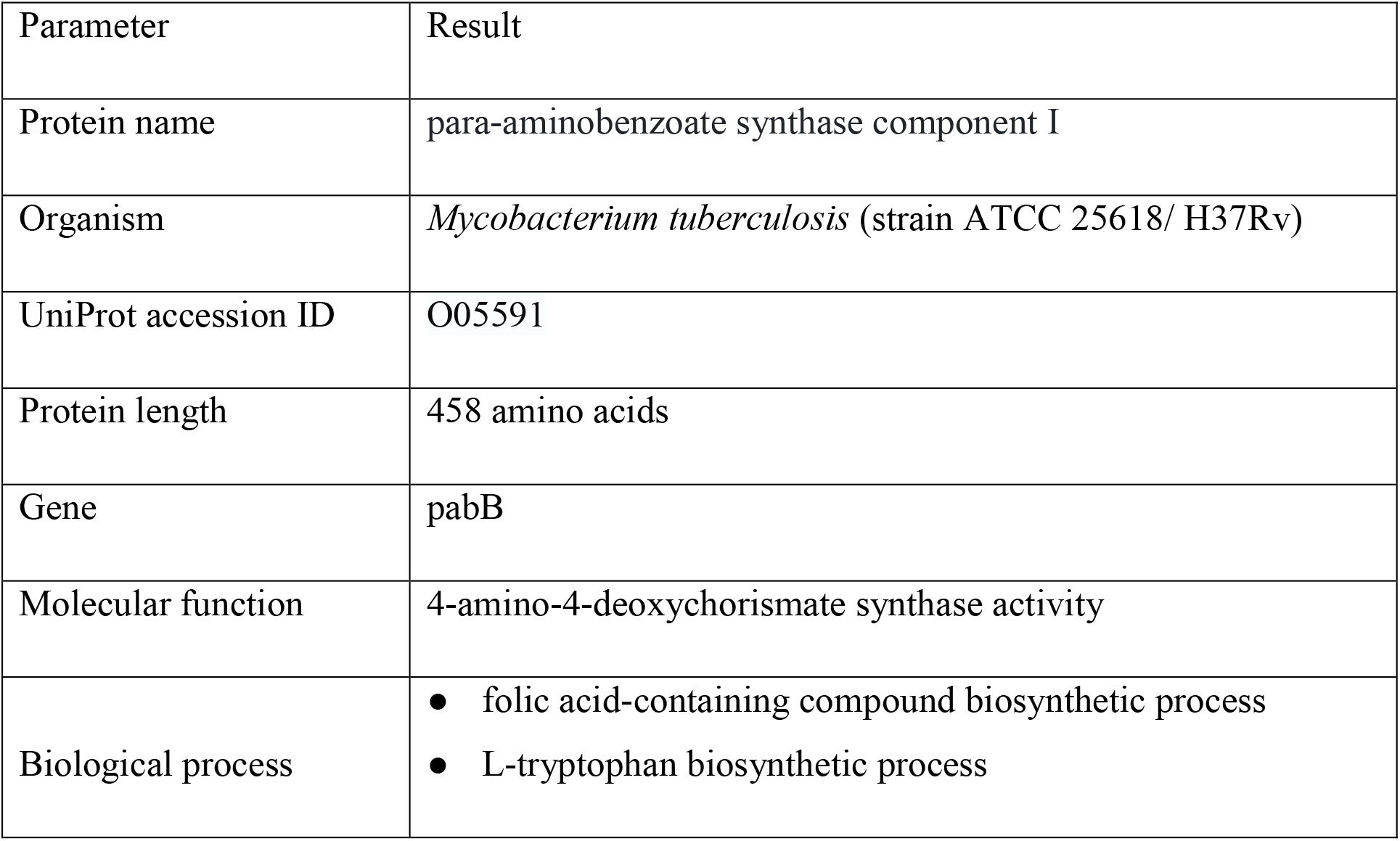
UniProtKB information of para-aminobenzoate synthase component I (PabB)

**Table 4.** KEGG Orthology(KO) identifiers of para-aminobenzoate synthase component I (PabB).

| Source | KO identifier(s) |
| --- | --- |
| UniProtKB | K01665 |
| DEG | K03342,K04781,K02552,K02361,K01657,K13950,<br>K01665 |

**Table 5.**
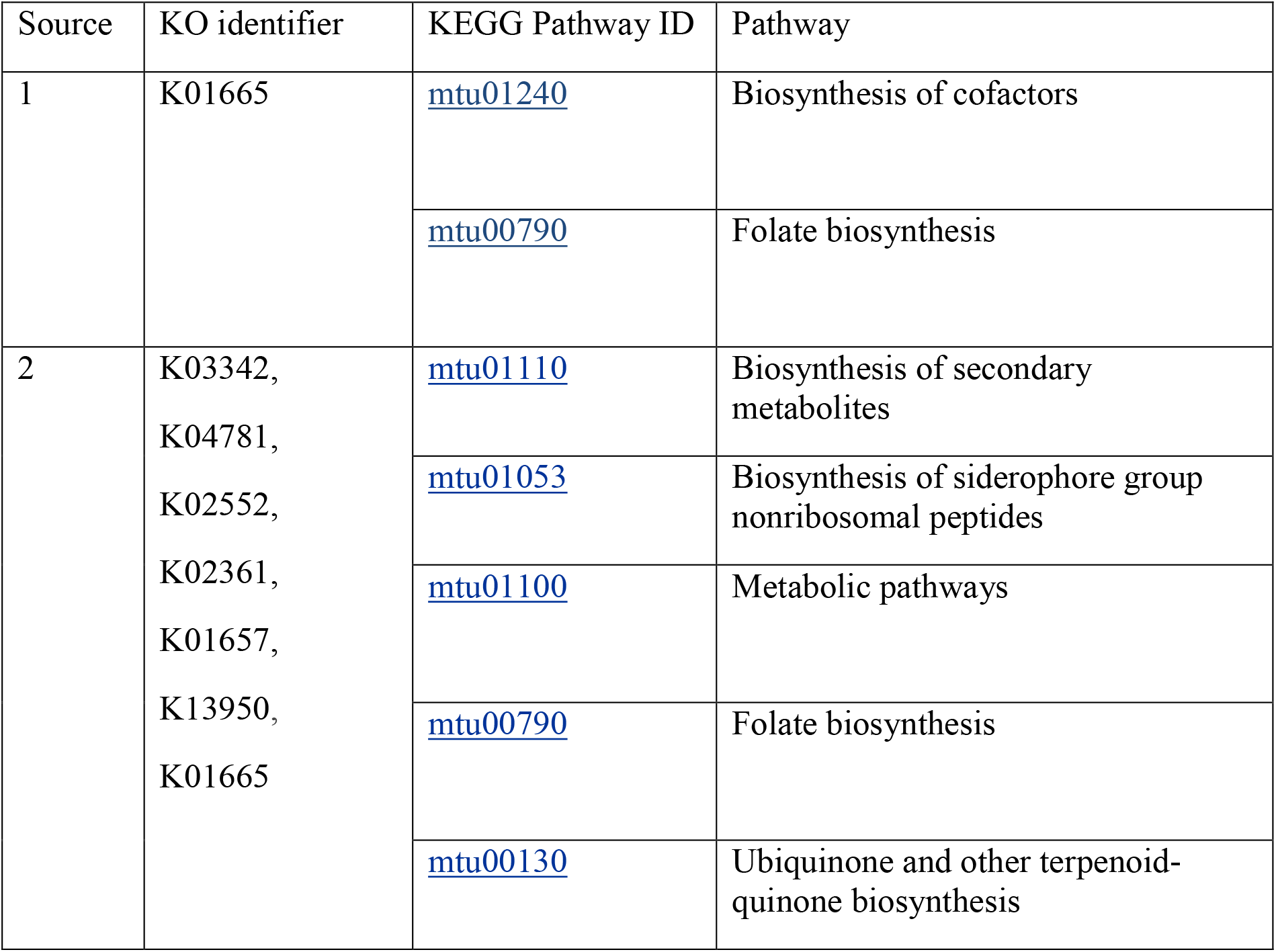
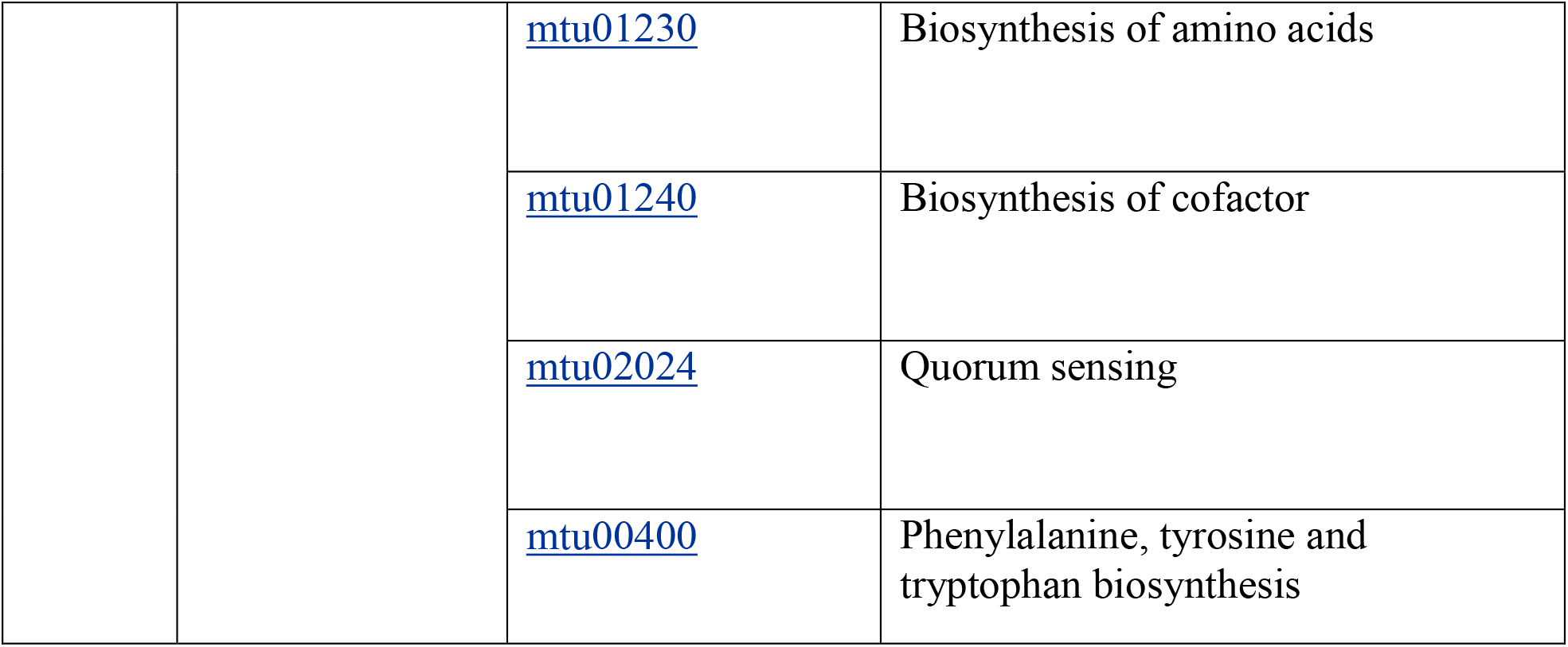
KO Identifier(s) and their corresponding pathways.

**Table 6.** Gene Ontology annotations of para-aminobenzoate synthase component I (PabB) obtained from QuickGO.

| GENE<br>PRODUCT ID | GENE<br>SYMBOL | GO TERM | GO NAME |
| --- | --- | --- | --- |
| O05591 | pabB | GO:0046820 | aminodeoxychorismate synthase activity |
| O05591 | pabB | GO:0046820 | aminodeoxychorismate synthase activity |
| O05591 | pabB | GO:0000162 | L-tryptophan biosynthetic process |
| O05591 | pabB | GO:0009396 | folic acid-containing compound<br>biosynthetic process |

**Table 7.** List of Top 10 drug candidate compounds prioritized from 59 ADMET-filtered compounds for molecular docking.

| Rank | Ligand No. | Catalog ID | SMILES | ADMET score | Compound name |
| --- | --- | --- | --- | --- | --- |
| 1 | Ligand 3 | Z1198155509 | CCC(C)C(CC)C(N)=O | 0.749776 | Valnoctamide |
| 2 | Ligand 27 | Z1198150037 | CN1CCC[C@H]1C2=CC=CN=C2 | 0.743602 | Nicotine |
| 3 | Ligand 13 | Z8173529772 | OC=1C=CC=CC1 | 0.729967 | Phenol |
| 4 | Ligand 1 | Z104473074 | CC1(C)C2CCC1(C)C(=O)C2 | 0.725247 | Camphor |
| 5 | Ligand 17 | Z104474028 | COC=1C=CC=CC1O | 0.722209 | Guaiacol |
| 6 | Ligand 6 | Z104476418 | OCC=1C=CC=CC1 | 0.720115 | Benzyl alcohol |
| 7 | Ligand 45 | Z1522567174 | Cl.NC12CC3CC(CC(C3)C2)C1 | 0.720023 | Amantadine Hydrochloride |
| 8 | Ligand 5 | Z104473570 | OCCOC=1C=CC=CC1 | 0.716794 | Phenoxyethanol |
| 9 | Ligand 48 | Z2238930991 | Cl.Cl.CNCCC=1C=CC=CN1 | 0.710086 | Betahistine hydrochloride |
| 10 | Ligand 20 | Z104473820 | CC=1C=CC=C(O)C1 | 0.708075 | M-Cresol |

**Table 8.** DiffDock docking confidence scores and GNINA validation scores of the selected ligands against PabB.

| Ligand | DiffDock Confidence | GNINA Minimized Affinity(kcal/mol) | GNINA CNN Score | GNINA CNN Affinity |
| --- | --- | --- | --- | --- |
| Camphor (Ligand 1) | -0.651958942 | -3.87263775 | 0.77934444 | 4.51715517 |
| Valnoctamide (Ligand 3) | -1.15039885 | -3.2714591 | 0.33411473 | 3.58859253 |
| Phenoxyethanol (Ligand 5) | -0.431559593 | -4.81692553 | 0.557757854 | 3.46202874 |
| Benzyl alcohol (Ligand 6) | -1.205140591 | -2.45232797 | 0.523474276 | 2.79231763 |
| Phenol (Ligand 13) | -0.369805604 | -2.40141129 | 0.905413806 | 4.35881853 |
| Guaiacol (Ligand 17) | -0.860946775 | -3.41519547 | 0.826025426 | 3.61435056 |
| M-Cresol<br>(Ligand 20) | -0.641652882 | -3.96926403 | 0.840731978 | 3.79526019 |
| Nicotine<br>(Ligand 27) | -0.0320611 | -4.38831997 | 0.666002274 | 4.44382477 |

**Table 9.** Interaction map of best docked conformations of ligands from GNINA.

| Docking conformation | Binding energy (kcal/mol) | Interacting residues (Hydrogen bonds) | Interacting residues (Hydrophobic interactions) |
| --- | --- | --- | --- |
| PabB vs. Camphor (Ligand 1) | -3.87263775 | SER252 | TRP68 , ARG239, VAL385,ASN397,ALA399,ILE400 |
| PabB vs. Phenoxyethanol (Ligand 5) | -4.81692553 | SER252, VAL385, ASN397 | GLU256, VAL385, ASN397, VAL398, ALA399, ILE400 |
| PabB vs. M-Cresol (Ligand 20) | -3.96926403 | ILE400 | ARG239, GLU256, VAL385, ASN397, VAL398 |
| PabB vs. Nicotine (Ligand 27) | -4.38831997 | SER252, ASN397, ALA399, ILE400 | ARG239, GLU256, VAL385, ASN397, VAL398, ALA399 |

**Figure 7.**
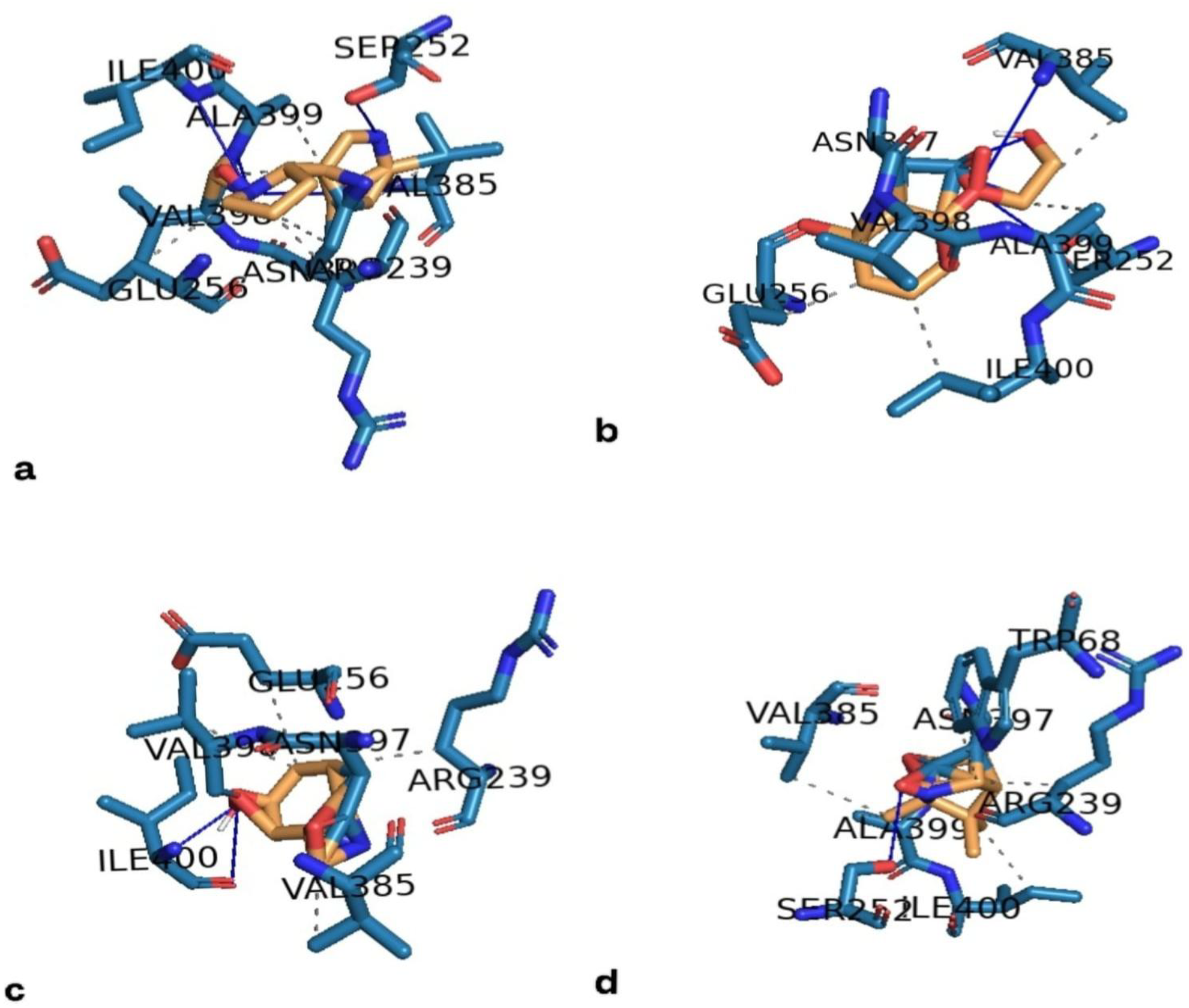
Docked Conformations of Camphor (Ligand 1), Phenoxyethanol (Ligand 5), M-Cresol (Ligand 20), Nicotine (Ligand 27) along with their interactions are shown in a,b,c,d respectively.

Among the analysed compounds, Phenoxyethanol (Ligand 5) and Nicotine(Ligand 27) demonstrated comparatively favourable docking energies and have multiple interactions in the docked conformations and therefore represent promising candidates for further investigation.

## Discussion

The present study was aimed at identifying potential drug targets in *M. tuberculosis* and the compounds targeting the same by using a computational approach. The study began with the screening of pathogen specific proteins involved in metabolic pathways crucial for survival of *M. tuberculosis* by using a pathway based analysis. The PabB protein involved in the folate biosynthesis pathway was prioritized and was used for further investigation. After PabB was chosen as a potential drug target, an FDA-approved drug library was screened based on ADMET properties and drug-likeness. The screened compounds were subjected to molecular docking against the protein using DiffDock. The docking results were validated using GNINA and the selected docking conformations showed hydrogen bond and hydrophobic interactions involving several residues in the predicted binding region after PLIP analysis. Among the screened compounds, Phenoxyethanol (Ligand 5) showed most favourable predicted binding energy (-4.82 kcal/mol) followed by Nicotine (Ligand 27) (-4.39 kcal/mol). Residues such as SER252, VAL385 and ASN397 were consistently involved in interactions across the analysed complexes, suggesting a common binding region for all the ligands within PabB.

Overall, the present study provides a computational approach for narrowing down the drug targets in *M. tuberculosis* and for investigation of potential compounds to be employed as drugs against the same. In future studies, the prioritized compounds can be used as a starting point to investigate their pharmacological properties and can serve as candidates for drug repurposing. However, the present findings are based on computational predictions, and experimental studies will be required to confirm their binding, inhibitory activity and anti-tubercular potential.

